# A Simple, Low-Cost, and Efficient Protocol for Rapid Isolation of Pathogenic Bacteria from Human Blood

**DOI:** 10.1101/2025.01.14.633023

**Authors:** Fatma S. Coskun, Joshua Quick, Erdal Toprak

## Abstract

Bacteremia is a serious clinical condition in which pathogenic bacteria enter the bloodstream, putting patients at risk of septic shock and necessitating antibiotic treatment. Choosing the most effective antibiotic is crucial not only for resolving the infection but also for minimizing side effects, such as dysbiosis in the healthy microbiome and reducing the selection pressure for antibiotic resistance. This requires prompt identification of the pathogen and antibiotic susceptibility testing, yet these processes are inherently slow in standard clinical microbiology labs due to reliance on growth-based assays. Although alternative methods exist, they are rarely adopted in clinical settings because they involve complex protocols and high costs for training and infrastructure. Here, we present an optimized, simple protocol for rapidly and efficiently isolating bacterial pathogens from blood without altering typical laboratory workflows. Our method is cost-effective and compatible with commonly available laboratory instruments, offering the advantage of isolating bacterial cells directly, which bypasses the delays associated with traditional blood culture methods and enables faster diagnostic results. The protocol achieved over 70% efficiency within 30 minutes, remained effective at low bacterial concentrations (1–10 bacteria/0.3 mL blood), and preserved bacterial viability with no notable change in growth lag times. We validated the protocol on several clinically relevant bacterial strains, including *Escherichia coli, Klebsiella pneumoniae*, and *Staphylococcus aureus*. These findings highlight our protocol’s potential utility in clinical and research settings, facilitating timely cultures and minimizing diagnostic delays.

## Introduction

Antibiotics are among the most important components of modern medicine, enabling clinicians to perform complex procedures such as surgeries, cancer treatments, and organ transplants. They are routinely used prophylactically and empirically to prevent and treat bacterial infections, and their introduction in the late 1920s has greatly increased life expectancy (1). However, the effectiveness of antibiotics has been significantly dampened by the rapid emergence of antibiotic resistance which limits their clinical use. According to the Centers for Disease Control and Prevention (CDC), antibiotic-resistant bacteria cause over 2.8 million infections and more than 35,000 deaths annually in the United States (2).

Pathogenic bacteria can translocate into the normally sterile bloodstream, originating either from external sources or the host microbiome, particularly in immunocompromised patients which then causes bacteremia. Treating bacteremia is clinically challenging due to various unknowns, such as the identity and quantity of the bacteria, their growth state, the presence of toxin-producing genes, and their antibiotic susceptibility profile. Without this information, clinicians must rely on empirical treatments to achieve favorable outcomes (3). As a result, patients—depending on their condition—are often immediately started on multiple broad-spectrum antibiotics to cover both Gram-negative and Gram-positive bacteria. Based on the patient’s response, some drugs may be discontinued while others are added. Although this approach can improve short-term outcomes, it also contributes to the global antibiotic resistance problem by selecting resistance-conferring mutations and genes in clinical settings.

The solution appears straightforward: clinicians should rapidly identify the bacteria causing the infection and determine their antibiotic susceptibility, allowing for evidence-based treatment regimens that minimize the use of unnecessary or ineffective antibiotics. However, this is a challenging task, as most common clinical microbiology tests are growth-based and therefore inherently slow. Blood drawn from the patient is incubated— either by plating or liquid culturing—until bacterial growth is detected, which can take several days (Figure 1A). Furthermore, it is not uncommon for attempts to culture pathogenic bacteria to fail, either due to prior antibiotic exposure or unsuitable laboratory conditions for bacterial growth. Another major technical limitation in diagnosing bacteremia is the crowded composition of human blood. Although genomics-based advanced sequencing and imaging techniques are available, their clinical utility remains limited. The genetic signals from abundant blood cells interfere with the diagnostic capacity of sequencing-based methods, while the high number of blood cells makes it extremely challenging to identify and localize bacterial cells. Therefore, a simple and robust tool that can isolate bacterial cells from blood without compromising bacterial viability would significantly enhance the effectiveness of these methods.

**Figure 1.**
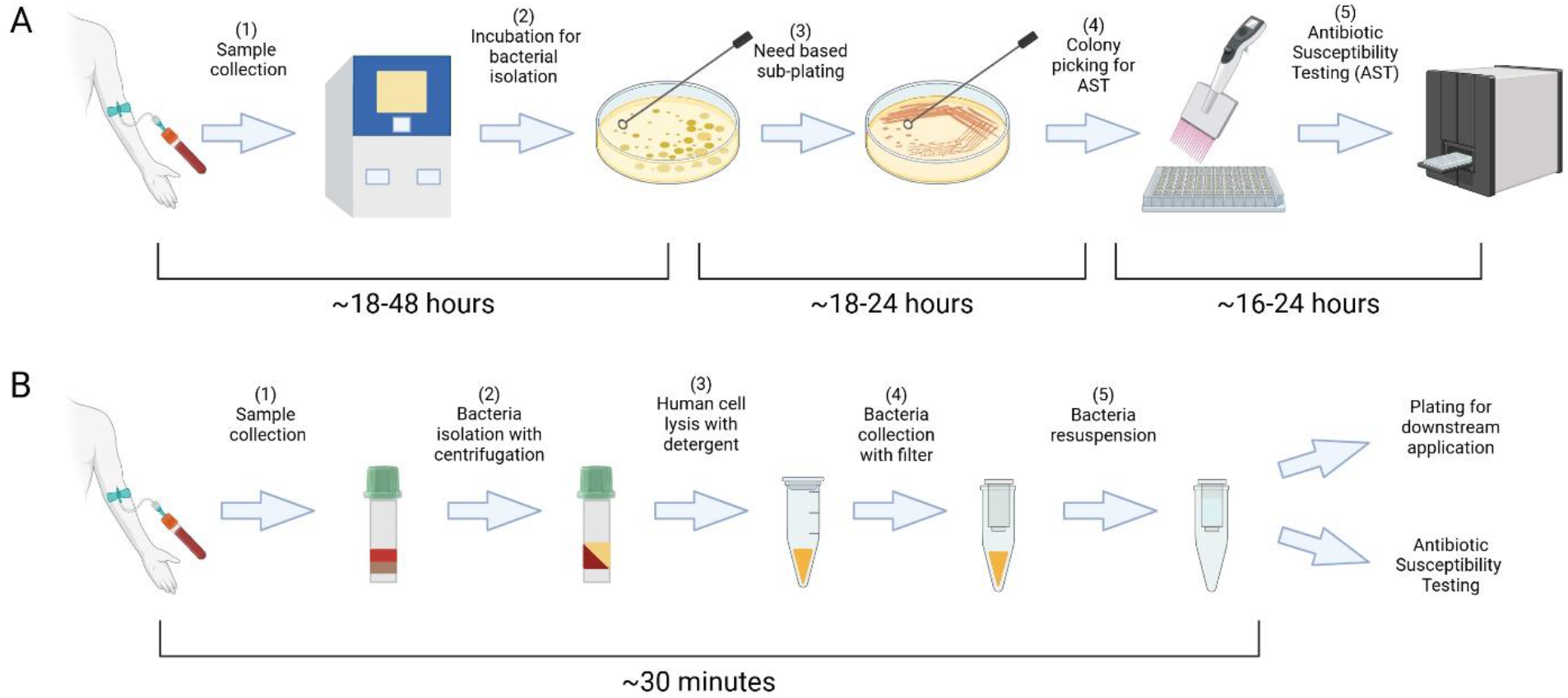
The standard protocol for managing infectious agents typically requires a period of 2 to 4 days for their isolation and characterization (A). Our enhanced protocol streamlines the bacterial isolation process, reducing the time required to just 30 minutes with readily available laboratory equipment and consumables (B).

Currently, the most common method for detecting bacteremia involves plating and incubating blood on special agar plates until bacterial colonies form. A more advanced approach uses diagnostic systems like BACTEC and BacT/ALERT. The BACTEC system detects bacterial growth by measuring ^14^CO_2_ release, which is produced as bacteria metabolize nutrients in the culture medium, making it highly sensitive to even small amounts of bacterial CO_2_ production (4). In contrast, BacT/ALERT uses colorimetric sensors to detect CO_2_ level changes, signaling microbial growth with minimal manual handling and reducing contamination risks (5, 6). Both systems automate the monitoring process, providing real-time alerts to laboratory personnel and enhancing detection speed and accuracy, thereby helping reduce treatment delays and healthcare costs by enabling targeted therapy. However, a significant limitation of these methods is their reliance on bacterial growth, which can take several days. Furthermore, blood cultures marked as positive for bacterial growth after long incubation times do not accurately reflect the true bacterial load in the patient’s blood; most cells in these samples are still blood cells, making them unsuitable for direct use with modern detection and phenotyping tools. When blood samples test positive in initial screenings, they undergo plating and even sub-plating for identification, introducing additional delays in the diagnostic process (Figure 1A).

Once disease-causing bacteria are isolated from blood as pure cultures or colonies, various conventional and novel techniques become available for downstream analysis, including bacterial identification, imaging, and antibiotic susceptibility testing (7, 8, 9, 10).

Standard antimicrobial testing methods include the disk diffusion assay and liquid-based assays, in which bacterial growth is quantified in multiple antibiotics at different doses. Traditional methods usually require one to three days, but newer approaches, like the Accelerate Pheno system that uses microfluidic approaches combined with fluorescence *in situ* hybridization (FISH) speed up the process with varying efficiency (11). Laser-based technologies like Raman spectroscopy enhanced by deep learning can differentiate bacterial species by analyzing their unique vibrational spectra (12). On the other hand, Matrix-Assisted Laser Desorption/Ionization Time-of-Flight Mass Spectrometry (MALDI-TOF MS) can identify microorganisms from pure cultures at the species level within minutes by profiling unique spectroscopic signatures (13, 14, 15, 16). In addition to these methods, rapid metagenomics using nanopore sequencing and rapid quantitative PCR are emerging as promising alternatives, enabling the detection of bacterial DNA or resistance genes in clinical samples (17, 18, 19, 20). Advanced technologies, such as the PA-100 AST nanofluidic system, which won the Antimicrobial Resistance (AMR) Longitude Prize, exemplify innovative solutions for rapid and precise AMR testing using minimal sample preparation. This system has undergone clinical evaluation, demonstrating its effectiveness in delivering phenotypic antimicrobial susceptibility test results within 45 minutes (21). In summary, all emerging techniques for identifying and phenotyping bacteria have shown promise for rapid and precise microbial detection. However, these advanced techniques often rely on or are significantly enhanced by the availability of pure bacterial cultures.

Techniques such as vacuum filtration (22), microfluidics-based approaches (23, 24), and flow-based systems (25) have been developed to enable direct pathogen identification from blood samples in clinical settings, bypassing the limitations of traditional culturing. Yet, a widely available method is still lacking for the direct isolation of bacteria from freshly drawn blood. To address this gap, we developed a simple, rapid, and low-cost protocol that reduces the time needed to obtain pure bacterial cultures from several days to just 30 minutes (Figure 1B). In brief, our protocol employs a detergent to selectively lyse host blood cells and uses filter centrifugation to isolate viable bacteria with over 70% efficiency. We successfully tested multiple clinically significant bacterial strains, including *Escherichia coli* (E. coli), *Klebsiella pneumoniae* (Kleb), and *Staphylococcus aureus* (S. aureus).

## Methods

### Bacterial strains

Initial optimization experiments were conducted using a clinical *E. coli* strain (ATEC) that was isolated from the feces of a pediatric stem cell transplant patient before any antibiotic treatment, as previously described (26). All other clinical strains, including *Escherichia coli* 3122 B34 (*E. coli*), *Klebsiella pneumoniae* 3160 B36 (*Kleb*), and *Staphylococcus aureus* 3220 B37 (*S. aureus*) were kindly provided by Dr. David Greenberg (UTSW). All bacterial strains were handled in strict adherence to biosafety level 2 (BSL2) regulations.

### Bacterial Spiking into Blood and Isolation

Overnight-grown bacterial cultures were normalized to an optical density of 1 at 600 nm (OD600, ∼5 × 10^8^ CFU/mL) and serially diluted in sterile 96-well plates. A predetermined number of bacterial cells, calculated based on initial OD600 vs. CFU calibrations, were spiked into freshly drawn blood collected using a K2 Vacutainer (BD367899). A 300 µL aliquot of the spiked blood was transferred to lithium-heparin blood separation tubes (BD365985) and centrifuged at 6000 × *g* for 2 minutes (Figure 1B, steps 1 and 2) to trap unwanted blood cells (e.g., red and white blood cells) in the heparin column. Plasma (∼100 µL) was carefully removed using a pipette, ensuring no contact with the gel, as bacteria were expected to be collected above the gel. To recover bacteria, 100 µL of PBS was added to the tube, the mixture was gently pipetted 2–3 times, and the entire sample was transferred to a sterile Eppendorf tube (Figure 1B, step 3). The top surface of the heparin column was washed with 100 µL PBS to maximize bacterial recovery. A 1% saponin solution (final concentration, Sigma-Aldrich SAE0073) was then added to the Eppendorf tube to lyse remaining blood cells, and the PBS-saponin mixture was vortexed for 15 seconds and incubated at room temperature for ∼15 minutes. The sample was subsequently transferred to a Spin-X column filter (0.2 µm membrane, Sigma-Aldrich CLS8162) and centrifuged at 6000 × *g* for 2 minutes (Figure 1B, step 4). After discarding the flow-through, bacterial cells trapped by the porous membrane were washed by adding 200 µL of PBS, vortexing for 15 seconds, and centrifuging again at 6000 × *g* for 2 minutes (Figure 1B, step 5). The flow-through was discarded, and bacteria were resuspended in 100 µL of sterile PBS by vortexing for 30 seconds. To increase recovery, the last step was occasionally repeated. Finally, the entire bacterial suspension was collected for further analysis. For viability testing, the bacterial suspension was evenly spread onto an agar plate using a sterile spreader loop, and colony-forming units (CFU) were enumerated the following day. As a control, a reference volume of bacterial cells was always plated prior to spiking bacteria into the blood to normalize CFU counts and calculate protocol efficiency. A more detailed protocol is shown in Figure S1.

### Quantifying Bacterial Viability with Lag Time Measurements

Lag time is typically defined as the duration before bacterial cells enter the exponential growth phase after being diluted to very low densities following overnight growth to saturation. In our experiments, we defined lag time as the time required for a bacterial culture to reach a background-corrected optical density (OD600) of 0.04, at which standard absorption-based methods are sensitive enough for detection and quantification. At this point, optical density measurements are no longer limited by the detection threshold of ∼0.005 in our plate reader (Tecan M200PRO). Bacterial cells (100 µL) isolated from blood were transferred to 96-well plates prefilled with 100 µL of 2x concentrated Luria Broth (LB) growth medium. Bacterial growth was monitored continuously by measuring OD600 every 3 minutes, and background-corrected OD600 readings were used to calculate the lag time. For each experiment, the lag time of bacteria not subjected to the isolation protocol was also measured to quantify its effect on bacterial growth delay. In parallel, the number of spiked and recovered viable bacteria was determined by plating cells on LB agar and counting colony-forming units (CFUs).

## Results

### We efficiently recover bacterial cells from human blood in 30 minutes

We developed and optimized a simple protocol for the rapid isolation of bacteria from human blood. This protocol involves basic centrifugation and pipetting steps, using standard laboratory equipment and commonly available consumables. Since it is challenging to access blood samples from patients suspected of bacteremia, particularly those who have not been treated with antibiotics, and because it is difficult to determine the exact number of bacteria per milliliter in such samples, we tested our protocol by spiking a known number of bacterial cells into freshly collected sterile human blood. Blood donors had not used any antibiotics for at least two weeks prior to blood collection.

As described in the methods section, a precalculated number of bacterial cells were spiked into sterile human blood to evaluate the efficiency of our protocol. As shown in Figure 2, we tested our protocol’s efficacy by spiking *E. coli* cells (ATEC, methods) with a range of cell densities into blood. Regardless of the number of bacteria spiked, we recovered more than 70% of the bacterial cells (Figure 2A). Remarkably, even when only one or two bacterial cells (1.8 ± 1.83) were spiked into the blood, we recovered most of the cells (1.2 ± 1.17). This result highlights the robustness of the protocol, even under conditions where stochastic noise due to low bacterial counts could complicate accurate cell counting.

**Figure 2.**
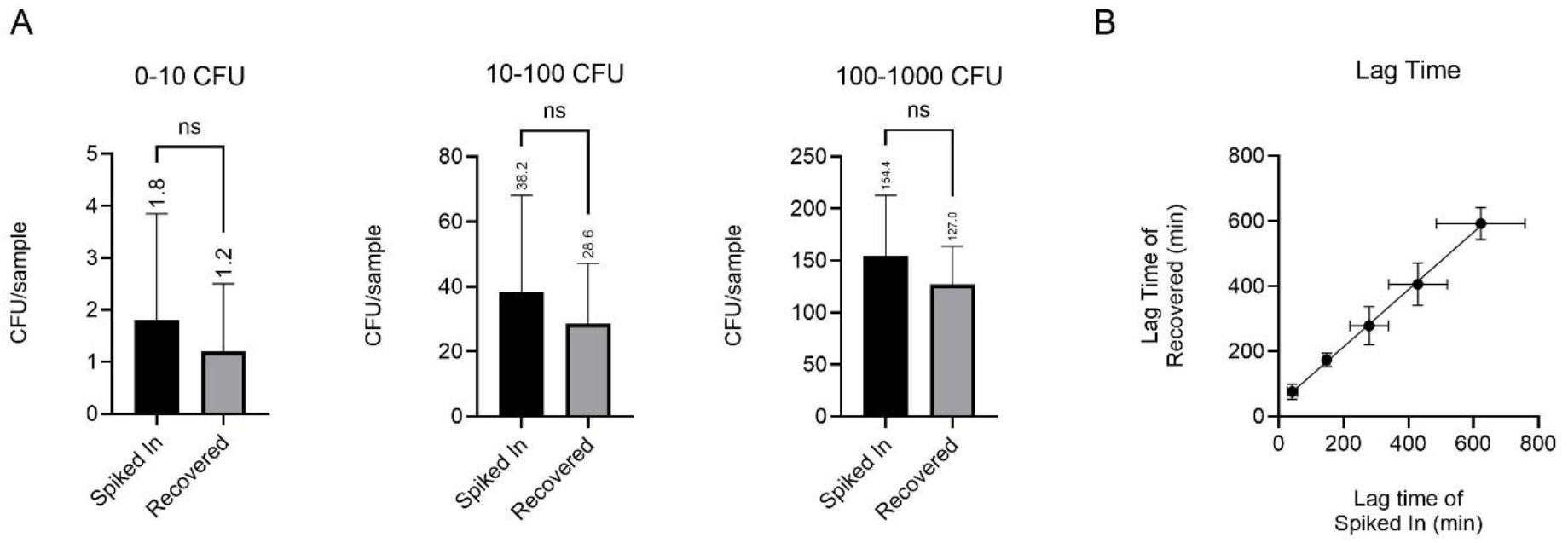
Our protocol achieved recovery rates exceeding 70%, even when tested with extremely low spiked densities of a clinical *E. coli* isolate (ATEC) (A). The lag time (in minutes) of the spiked-in and recovered ATEC cells is shown. The solid black line represents y = x with R^2^=0.9661) (B).

A valid concern regarding our bacterial isolation protocol is whether the detergent and washing steps introduce growth delays in recovered bacteria. To address this, we measured the lag times of spiked bacteria and compared them to those recovered from blood. As shown in Figure 2B, there were no prominent qualitative differences in lag times between the two groups (R^2^=0.9661). The only exception occurred at the lowest cell density tested, where stochastic noise, likely due to pipetting or other uncontrollable experimental factors such as phenotypic states of bacterial cells, had a measurable effect. In these cases, the lag time for recovered bacteria was approximately 30 minutes longer (roughly one doubling time) than that for bacteria not subjected to the isolation protocol. Otherwise, our protocol did not cause significant delays in bacterial growth. These recovered bacterial cultures can be directly used for identification or antibiotic susceptibility testing, as described previously. However, for sequencing purposes, an additional step involving nuclease treatment and subsequent washing may be required to eliminate host DNA or RNA contamination.

Upon optimizing the protocol with ATEC, we tested its efficacy against different bacterial species. For this, we used another clinical strain of *Escherichia coli* 3122 B34 (*E. coli*) and clinical strains of *Klebsiella pneumoniae* 3160 B36 (*Kleb*), as well as the Gram-positive *Staphylococcus aureus* 3220 B37 (*S. aureus*). All these species are considered pathogens of concern according to CDC reports. Our protocol demonstrated high efficiency across a range of pathogens, as shown in Figure 3, with recovery rates of approximately 90% for Gram-negative bacteria and around 50% for Gram-positive bacteria. The lower recovery efficiency for Gram-positive bacteria is likely due to their distinct morphological characteristics. Despite variations in growth characteristics and metabolic requirements among these bacteria, the protocol proved highly versatile, preserving bacterial viability effectively for a wide spectrum of pathogens commonly linked to bloodstream infections. Therefore, we expect this protocol to have broad utility in clinical microbiology laboratories.

**Figure 3.**
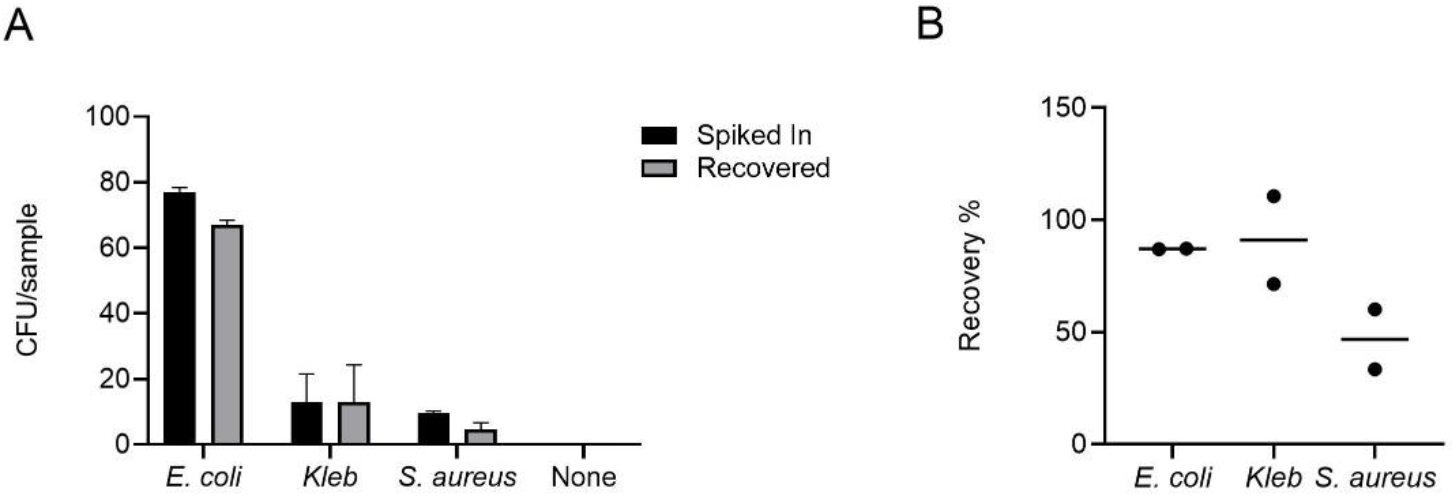
Our protocol demonstrated high efficiency across a diverse array of clinical isolates, encompassing both Gram-positive and Gram-negative bacterial species (A). The recovery rates, expressed as percentiles, achieved using this method are shown. (B).

## Discussion

Antibiotic resistance is a critical public health concern, and any intervention that significantly addresses this challenge and improves health outcomes is highly valuable.

Rapid bacterial detection, identification, and antibiotic susceptibility testing are essential for enabling clinicians to make evidence-based decisions when designing antibiotic treatment regimens. Despite numerous advancements in state-of-the-art methods over the past decade, their widespread adoption in clinical settings remains limited. This is largely due to high costs, the need for specialized infrastructure, and the additional burden of retraining clinical microbiology personnel, who are often already overextended.

There is an urgent need for simple, cost-effective methodologies that can expedite pathogen detection, identification, and phenotyping without disrupting the workflows of clinical microbiology laboratories. Our protocol is specifically designed to utilize commonly available lab instruments and consumables, making it practical for resource-constrained settings while integrating seamlessly into existing workflows By significantly reducing the time required for bacterial isolation, our method can integrate seamlessly with clinical workflows, allowing faster access to pure bacterial cultures for downstream diagnostic technologies, thereby enabling evidence-based decisions on antibiotic treatment and improving patient outcomes.

While accelerating bacterial isolation, our protocol primarily serves as a foundational step, providing pure bacterial cultures free of large host cells and preserving high bacterial viability, which are crucial for the effectiveness of state-of-the-art detection and phenotyping technologies. This adaptability enhances its potential for seamless integration into routine clinical workflows. Additionally, modified versions of our protocol could be developed for isolating specific pathogens, such as fungal cells, from the bloodstream. Given these advantages, we anticipate that our methods will generate broad interest and widespread adoption among clinical microbiologists.

## Acknowledgements

This study was supported by the Advanced Research Projects Agency for Health (ARPA-H) (1AYSAX000005-01), NIH grant R01GM125748, UTSW High Risk High Impact Award, and Welch Foundation I-2082-20240404. We thank Dr. Fatih Cansizoglu for his contributions to the optimization of the bacteria isolation protocol.

