## Supplementary Figure 1 for "A Simple, Low-Cost, and Efficient Protocol for Rapid Isolation of Pathogenic Bacteria from Human Blood"

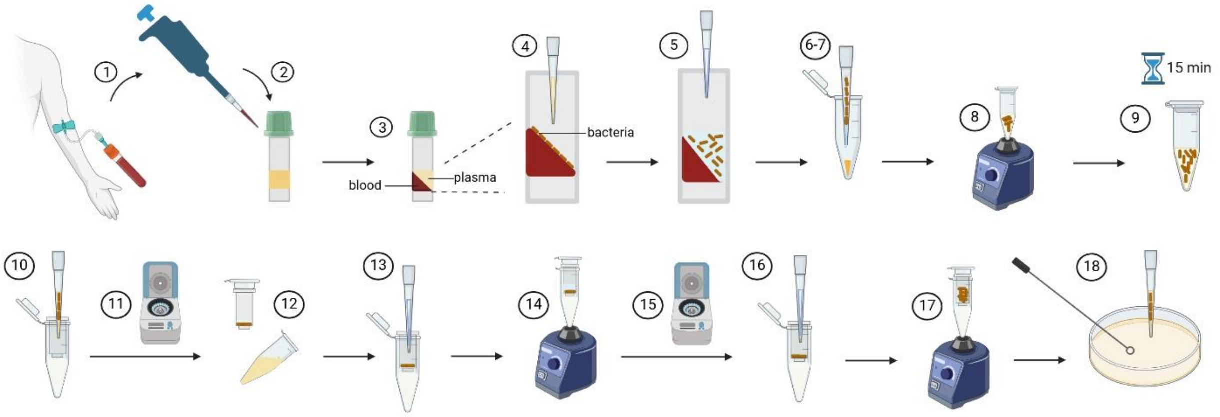


**Figure S1.** 1) 5 ml of blood is collected in a K2 Vacutainer. 2) 500 µl of blood is transferred to a serum separation tube (SST). 3) The SST is centrifuged at 6000 g for 2 minutes. 4) The plasma is carefully removed, avoiding contact with the gel, as bacteria will be on the gel's surface. 5) 100 µl of PBS is added to the gel's surface and pipetted gently 2–3 times. 6) The mixture is transferred to an Eppendorf tube containing saponin. 7) Steps 5 and 6 are repeated. 8) The sample is vortexed for 15 seconds. 9) The Eppendorf tube is incubated at room temperature for 15 minutes. 10) The sample is transferred to a spin-X column. 11) The spin-X column is centrifuged at 6000 g for 2 minutes. 12) The filter is removed, the flow-through is discarded, and it is placed back into the tube. 13) The filter is washed by adding 200 µl of PBS to its surface. 14) It is vortexed for 15 seconds. 15) It is centrifuged at 6000 g for 2 minutes and the flow-through is discarded. 16) To collect the bacteria, 100 µl of PBS is added to the filter's surface. 17) It is vortexed for 30 seconds. 18) The liquid from the filter's surface is transferred to a plate and spread evenly.
